# Arabidopsis Membrane Contact Site protein SYNAPTOTAGMIN A maintains sieve element endomembrane morphology and function

**DOI:** 10.64898/2026.01.26.701828

**Authors:** Chiara Bernardini, Stacy Welker, Christopher Vincent, Maryam Khalilzadeh, Rita Musetti, Aart J E Van Bel, Amit Levy

**Affiliations:** Citrus Research and Education Center, University of Florida, Lake Alfred, FL, 33850, USA Department of Agriculture, Food, Animal and Environment, Università degli Studi di Udine, Udine, Italy; Department of Agriculture, Food, Animal and Environment, Università degli Studi di Udine, Udine, Italy; Horticultural Sciences Department, University of Florida, Gainesville, FL 32611; Department of Land, Environment, Agriculture and Forestry (TESAF), Università di Padova, via dell’ Università, 16, 35020 Legnaro, PD, Italy; Institute of Phytopathology, Justus-Liebig University, Heinrich-Buff-Ring 26–32, 35392 Giessen, Germany; Department of Plant Pathology, University of Florida, Gainesville, FL 32611

**Keywords:** Synaptotagmin, Membrane contact sites, Sieve element endoplasmic reticulum, Phloem, Phloem infection

## Abstract

Very little is known about the sieve element (SE) endomembrane system. In terms of surface area, the most important membrane is the SE endoplasmic reticulum, characterized by a unique cisternal structure of anchored flat and smooth stacks. Plasma membrane contact sites (MCSs) play a crucial role in anchoring the endoplasmic reticulum (ER) to the plasma membrane (PM) and in maintaining membrane intergrity. Here, we tested their role of MCSs in the endomembrane system of the sieve elements. Synaptotagmin A (SYTA) is one of the best-studied proteins known to form ER-PM MCSs in plants. We show that SYTA:RFP co-localizes with SUC2::GFP and CALS7::GFP, confirming its presence in *Arabidopsis* SEs. In *syta-1* mutants, SER lost its discrete shape and separated from the SE wall. The export of ^14^C-compounds from leaves of *wild type* plants was about 10% higher than in *syta-1* mutants. Finally, we explored SYTA’s role in phytoplasma infection response. After infection, *syta-1* plants displayed 50% less callose deposition and an uneven distribution pattern of the pathogen. In conclusion, our work shows that SYTA is required for maintaining the unique shape of the SE endomembrane system, and for diverse SE functions including callose deposition, carbon translocation and response to pathogens.

## Introduction

The sieve-element endomembrane system is unique among the plant membrane structures, and its functions are poorly understood. The sieve element ER (SER) is composed of stacks of cisternae without ribosomes, arranged either parallel or perpendicular to the plasma membrane (PM) and evolves from a rough reticulum to a smooth one during sieve-element (SE) maturation at the time that the nucleus degenerates (Behnke and Sjolund, 2012). The SER was reported to be sometimes associated with a mitochondrion (Sjolund and Shih, 1983). A later study demonstrated that SER is consistently connected to mitochondria and P-plastids (Ehlers et al., 2000). For efficient translocation through mature SEs, it is essential that the SER remains parietal and pushed against the cell wall (Musetti et al., 2023; Thorsch and Esau, 1981). Hence, SER has a clamp-like anchoring system that helps maintain its parietal stability and prevents it from being dragged along with the mass flow (Ehlers et al., 2000). The first type of clamps serves as an anchoring system between SER and other organelles or SER and the PM. A second type of clamps could occur between SER stacks and could provide the key in understanding the SER structure (Ehlers et al., 2000). However, the identity of these anchors remains unknown and their molecular structure is anticipated to be diverse due to the variable anchorages.

Potential tools for membrane anchoring may be provided by membrane contact sites (MCSs), which are specialized regions where two membranes come into close proximity, typically within 10 nm, excluding ribosomes (Baillie et al., 2020; Voeltz et al., 2024; Wang et al., 2017). These sites facilitate direct, non-vesicular movement of small molecules between organelles, mediated by shuttle proteins or protein complexes, and are involved in tethering organelles to the plasma membrane or the membrane of other organelles (Prinz et al., 2020; Voeltz et al., 2024). Due to their diverse protein composition (Pan et al., 2024), MCSs perform a variety of essential functions, including anchoring the highly dynamic organelle network, monitoring phospholipid metabolism and signaling, and facilitating lipid synthesis and transport pathways (Wang et al., 2017). ER-plasma membrane contact sites (EPCSs) generate attachment sites between the ER and the PM that facilitate Ca^2+^-dependent lipid exchange, Ca^2+^ homeostasis, non-vesicular lipid transfer, and membrane resealing (Pérez-Sancho et al., 2015; Schapire et al., 2008; Voeltz et al., 2024; Yamazaki et al., 2008). EPCSs also function at plasmodesmata, where the desmotubules (ER-derived membrane tubules between adjacent cells) are attached to the PM and where they are postulated to regulate plasmodesmal gating. Alterations in PD configuration are thought to be associated with modifications of the EPCSs (Bayer and Benitez-Alfonso, 2024; Wang et al., 2017).

Plant synaptotagmins (SYTs) are parts of EPCSs and are orthologous to mammalian Extended synaptotagmins (E-syts) (Craxton, 2010; Yamazaki et al., 2010). In Arabidopsis, the genome encodes six synaptotagmins, including the recently identified *SYT7*. E-Syts share structural similarities with yeast tricalbins (Tcbs), including multiple C2 domains, an N-terminus transmembrane domain, and a synaptotagmin-like mitochondrial-lipid binding protein domain (SMP) (Toulmay and Prinz, 2012). However, plant SYTs diverge from E-SYTs and tricalbins in two key aspects: they possess only two C2 domains and attach to the membrane via the C2B domain rather than a hairpin structure (Juan L Benavente et al., 2021; Benitez-Fuente and Botella, 2023). Still, both plant synaptotagmins and the Extended synaptotagmins share a similar function in attaching to the ER membrane with their N-terminal transmembrane domains, and generating EPCs through the attachment of their C2 domains with the plasma membrane in a calcium-dependent manner (Benitez-Fuente and Botella, 2023). SYT-1 localizes in the immobile area of the ER, and knockdown of SYT-1 in Arabidopsis led to ER destabilization and the loss of polygonal ER structure (Siao et al., 2016; Ishikawa et al., 2018; Levy et al., 2015).

The synaptotagmins in *Arabidopsis* are organized in two big clusters, the SYT1-like and the SYT5-like, which differentiated early in the evolution of land plants (Ishikawa et al., 2020). These clusters mainly differ in the Ca^2+^ affinity of their C2B domain (García-Hernández et al., 2025). The protein SYT1 (hereafter referred to as SYTA) is the most extensively studied synaptotagmin. Besides its role at EPCSs, SYTA functions as a signaling intermediate during mechanical and abiotic stresses, such as osmotic imbalance or freezing (García-Hernández et al., 2025; Pérez-Sancho et al., 2015; Yamazaki et al., 2008). In this scenario, SYTA is responsible for the membrane stability maintenance: the association of the downregulation of SYTA and any kind of abiotic stress can lead to a lack of stability or, more severely, to the plasmolysis of the membrane (García-Hernández et al., 2025).

Similar to other family members (Wang et al., 2025), SYTA also plays a role in biotic stress responses, such as those induced by powdery mildew and viral infections (Kim et al., 2016; Levy and Tilsner, 2020). In Arabidopsis and *Nicotiana benthamiana*, SYTA interacts with viral movement proteins, facilitating virus translocation through the plasmodesmata and forming viral replication complexes (VRCs) near the ER (Levy et al., 2015; Lewis and Lazarowitz, 2010). SYTA is a promoter of viral infection: its downregulation delays the infection and movement of several virus genera (Lewis and Lazarowitz, 2010; Uchiyama et al., 2014), and reduces the damage induced by the virus (Tian et al., 2024). SYT4 has been shown to reside in the phloem (Kumar et al., 2024), and preliminary evidence suggests phloem localization for other family members (Kumar et al., 2024), but the role of synaptotagmins in the SEs remains poorly understood.

In this study, we aimed to investigate the role of SYTA and EPCSs in the configuration and function of the SE endomembrane system, focusing on the SER structure. Our findings demonstrate that SYTA is present in SEs, where it plays a key role in regulating the architecture of endomembrane system and in regulating carbon export. Additionally, we showed that the knockdown of SYTA affects SE-phytoplasma interaction by altering callose metabolism and modifying the distribution pattern of the pathogen. These results provide new insight into the mechanisms underlying the unique architecture of the SE endomembrane system and reveal a role of MCSs in SE biology.

## Material and methods

### Growth of Arabidopsis

Arabidopsis seeds (background Col-0) of *syta-1* (with AtSYTA downregulated) and syta-1;gSYTA-RFP line (expressing the promoter and genomic sequence of SYTA fused with a red fluorescent protein -RFP-in *syta-1* background) were previously reported by Levy et al. 2015. The use of SUC2::GFP and CALS7::GFP were previously described (Imlau et al., 1999; Kalmbach et al., 2023). The *wild type* line (ecotype Col-0) was used as the control. Plants were grown on Murashige and Skoog agar medium (4.3g MS salts, 1ml Gamborg’s vitamin 1000X, 10g sucrose, 0.5g MES, and 0.8% agar per liter) at 22°C under short day conditions (10 hours light/14 hours dark). The syta-1 line contains the gene for BASTA resistance, thus for the SYTA line screening, 2ug/ml of Bialaphos (BASTA-containing commercial product, Goldbio, St Luis, MO, USA) was added to the substrate. After 15 days, plants were transferred to a soil mixture for further growth.

### GUS assays

The GUS (β-glucuronidase) assay was performed to assess the localization of SYTA in the Arabidopsis tissues. Arabidopsis leaf disks were incubated in GUS staining buffer (1 mM X-Gluc, 50 mM phosphate-buffered saline (PBS) at pH 6.8, 20% v/v methanol, and 1% v/v Triton X-100) for 24 hours at room temperature in the dark. Leaf disks were then bleached with a solution of ethanol and acetone (4:1) for 48 hours at 65°C to remove the chlorophyll.

### Arabidopsis crossing and SYTA localization

To assess the presence of the SYTA inside SEs, we examined mutant plants where a phloem-residing protein, either SUC2 or CALS7, was fused with GFP. These lines were crossed with a *syta-1*;gSYTA-RFP line, in which SYTA was fused with RFP. We then observed colocalization of the fluorescent proteins.

Crossing was performed as reported by The Arabidopsis Information Resource -TAIR- (Berardini et al., 2015). Briefly, the anthers were removed and the emasculated inflorescence was pollinated from a mature flower of the other line. Siliques were harvested three weeks after crossing, and the seeds were subsequently grown on selective substrates, as described above. From the crosses *syta-1*;gSYTA-RFP x SUC2::GFP, and *syta-1*;gSYTA-RFP x CALS7::GFP plants were obtained. Roots from 15-day-old plants of wild type, and *syta-1*;gSYTA-RFP and the two crosses i.e. *syta-1*;gSYTA-RFP x SUC2::GFP, and *syta-1*;gSYTA-RFP x CALS7::GFP were examined using a Leica SP8 LSCM (Leica Microsystems Inc., Buffalo Grove, IL, USA) equipped with a 40x oil immersion objective. RFP and GFP localizations were visualized using excitation laser lines of 488nm. Emissions were detected at 561-633nm and 490-520nm, respectively. *wild type* plants were used to detect traces of autofluorescence.

### TEM observation and basic sieve element parameters

To examine and describe the SER structure, a transmission electron microscopy analysis was performed following the standard procedure described in previous studies (Bernardini et al., 2022b; Folimonova and Achor, 2010). Fully developed leaf main veins were fixed with 3% (v/v) glutaraldehyde in a solution 0.1 M of potassium phosphate buffer (pH 7.2) for 4 hours at room temperature. After fixation, the samples were washed in phosphate buffer and postfixed in 2% osmium tetroxide (w/v) in the same buffer for 4 hours at room temperature. The samples were further washed in the phosphate buffer, dehydrated through a series of 10% to 100% acetone (v/v) steps (increasing by 10% and waiting for 10 minutes per step), and gradually infiltrated and embedded in Spurr’s resin over a 3-day period. The fully infiltrated samples were baked in an oven at 70°C overnight. Ultrathin sections (thickness of 100 nm) were obtained with a Reichert-Jung Ultracut E microtome and mounted onto 200-mesh formvar-coated copper grids. Sections were stained with 2% aqueous uranyl acetate (w/v) followed by Reynolds lead citrate and examined with a Morgagni 268 transmission electron microscope (FEI). For image analysis, FIJI software (Schindelin et al., 2012) was used to quantify the number of SEs exhibiting abnormalities, the cross-sectional areas of SEs, the number of vesicles per SE, and the state of SER-PM attachment. The diameter of well-formed vesicles in the SEs was also measured.

### Ilastik process and preparation of the final mask

To create a particles’ mask of the ER and plasma membrane, we used both FIJI and Ilastik (Berg et al., 2019) software. A brief description of the methodology used to obtain the mask is presented in Figure 1. We normalized the pixels of the pictures using a Gaussian blur with a radius of 1.5 pixels with FIJI.

**Figure 1.**
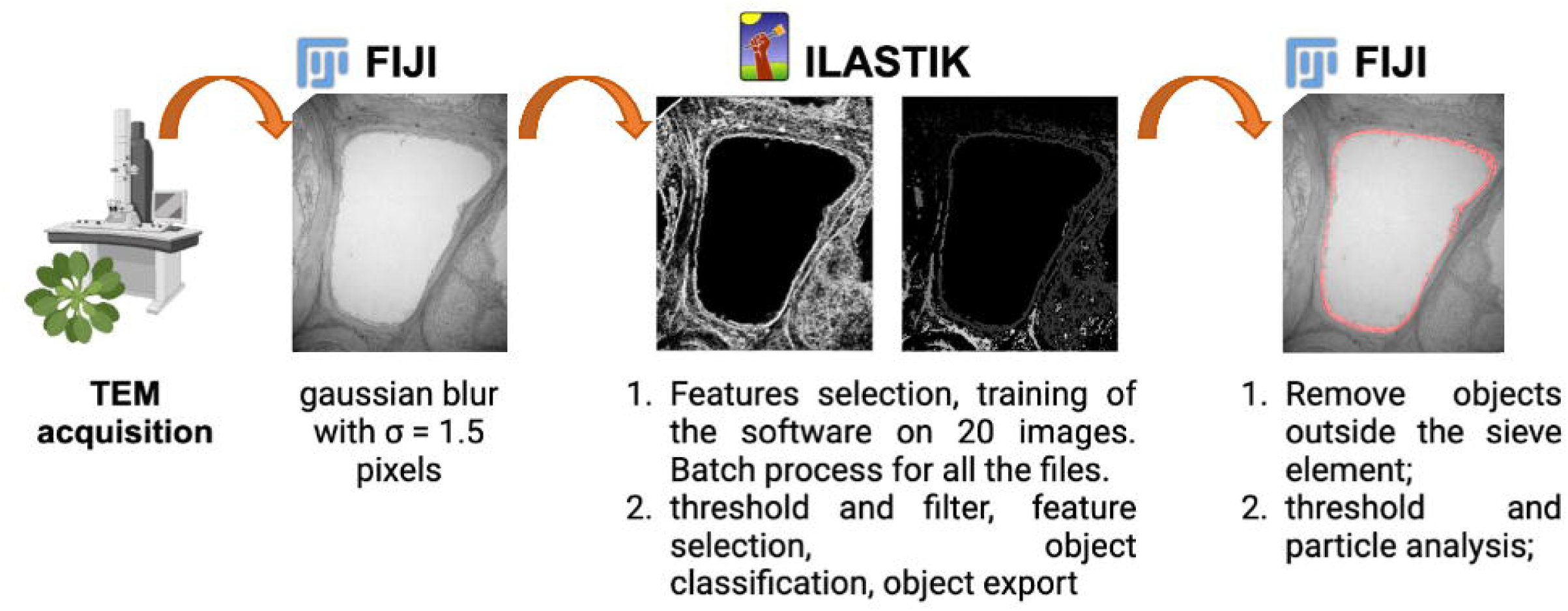
Workflow for creating an object prediction mask. Pictures were obtained with TEM. A Gaussian blur of the whole picture with sigma 1.5 was used to equalize the pixels used for the analysis in FIJI software. The obtained tiff images were used to train Ilastik software. 20 pictures (randomly picked from each treatment) were used as training for the machine learning procedure, and then the process was run in batch on all the files. Non-ER/PM objects were manually removed with the FIJI software before running the particle analysis on the ER pixels.

Using a supervised machine learning process, we trained the Ilastik software to distinguish SER-PM pixels from those of the SE lumen. A total of twenty images (randomly picked from each treatment) were used for the training process that involved all the available features for color and intensity, resemblance to an edge, and texture from σ0 to σ6. A Gaussian smoothing with σ=0.3 was used in the creation of the mask. The script was run in batch mode for the entire set of images to obtain probability mask .h5 files. Masks and original pictures were used for the object classification workflow in Ilastik. The probability mask was filtered for the layers of the ER and plasma membrane, and a simple threshold with a value of 0.5 was applied. The final object identity outputs were exported as .tiff files that were imported into FIJI and scaled to microns. Every object outside the SE was manually removed before the final object identity file was saved for pixellar particle analysis. A manual local threshold of 5-255 was applied to analyze the particles. Thus, every pixel with color values between 5 and up to 255 was considered as SER. The requested parameters were area, shape descriptors, skewness, kurtosis, Feret diameter and perimeter. The macro to process all the images is as follows:

dir1 = getDirectory(“Get Directory”);
list = getFileList(dir1);
setBatchMode(true);
for (i = 0; i < list.length; i++) {
showProgress(i + 1, list.length);
open(dir1 + list[i]);
run(“8-bit”);
run(“Manual Threshold…”, “min=5 max=255”);
run(“Analyze Particles…”, “display exclude summarize”);}
saveAs(“Results”, “/Users/chiarabernardini/Desktop/ER-segmentation/gaussian_blur3_ok/ER_filter/Summary.csv”);
close()

For our study, the aggregate SER-PM area expressed as μm^2^ per cell, the aggregate SER-PM perimeters expressed as μm, the Feret diameter expressed as nm, and the skewness and kurtosis expressed as arbitrary units were taken into consideration as crucial parameters. The Feret diameter represents the distance between the edge of the SER membrane border and the center of the shape, the skewness expresses the symmetry of the shape, and the kurtosis describes the flatness of the shape. For the entire analysis, at least 29 non-serial cells from 3 biological replicates per line were observed and analyzed.

### 14C export from leaves

We followed an established protocol to evaluate the photoassimilate export of *wild type* and *syta-1* lines (Bernardini et al., 2022a; Vincent et al., 2019). Briefly, Arabidopsis plants were grown, as reported above, under long-day light conditions (14 h L/10 h D) at 23°C. Sixty-day-old healthy plants of both lines were used for measurement. An x-ray photomultiplier tube, 5.6 cm in diameter (St. Gobain, Malvern, PA, USA), was used to detect the x-ray derived from the interaction of the ^14^C-emitted β-particles with the surrounding environment in the rosette leaves, from a distance of roughly 2-3 cm. The detector was connected to an M4612 12-channel counter, and the X-ray counts were logged using the manufacturer’s software (Ludlum Measurements, Sweetwater, TX, USA). Plants were placed on a layer of lead with a disposable 7ml weighing tray used as a receptacle under the same light conditions for the Arabidopsis growth (10 hours light/14 hours dark). Plants were then sealed in a plastic bag. After at least 8 hours of background measurement, 600μl of NaH^14^CO_3_ solution (specific activity: 40–60 mCi (1.48–2.22 GBq)/ mmol) were inject through a needle and volatized by adding 1000μl of 1M citric acid solution. After a 2-hour assimilation period, the bag was opened, and the gaseous radiolabeled isotope was allowed to evaporate out via a dedicated fume hood. The X-ray detector monitored the X-ray counts from the plant rosette every minute for 72 hours in the same light conditions. For each line, at least 3 biological replicates were carried out.

The data were processed using RStudio software (RStudio Team (2020). RStudio: Integrated Development for R. RStudio, PBC, Boston, MA). The counts collected from the channel counter were standardized from 0 to 1 to allow the comparison between different replicates. The standardized value (stV) was calculated as follows:

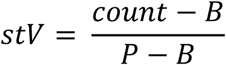

wherein B represents the background level taken before the isotope injection begins, and P is the average value at the peak of the curve (end of the injection).

We analyzed the DPMs and the stVs following two different workflows:

1. The general export curve during the time of experiment,
2. The loss of radioactive isotope from the rosette during the exponential phase described by a decay function.

The course of the curve was analyzed by selecting 12 key time points during the experiment (0, 250, 500, 750, 1000, 1250, 1500, 1750, 2000, 2250, 2500, and 2750 mins). The average value for each point was calculated and plotted for both the standardized and DPM values.

We filtered the stVs data from the starting point of the exponential phase of the curve to the end and generated its mathematical function. A self-start decay function was fitted to the data, and all the parameters of the curve were determined.

### Microscopic observations on phytoplasma distribution in *AtsytA* mutants

Seeds of *wild type* and *syta-1* were grown as described before (Bernardini et al., 2020). 50-day-old Arabidopsis plants from both the *syta-1* line and *wild type* lines were infected with Chrysanthemum yellow phytoplasma (group 16S-IrB) using the vector insect *Macrosteles quadripunctulatus* following a reliable protocol (Bernardini et al., 2022a). For each line, 10 plants were treated with infected vector insects, while 10 plants were treated with healthy insects of the same age and used as a control. The inoculation period for this experiment lasted 1 week. After inoculation, chlorophyll content was evaluated by measuring the light transmittance of three fully expanded leaves from each plant using a portable chlorophyll meter SPAD-502 (Minolta, Osaka, Japan), as previously reported (Pagliari et al., 2017), and the results were presented as SPAD index values. Data were taken weekly until the plants were harvested and their fresh weight determined. For TEM analysis, midribs of mature leaves were excised from the lamina and sectioned into 1 cm-long pieces and embedded following the standard procedure previously reported (Bernardini et al., 2020).

For the callose evaluation, midrib segments of mature leaves (1 cm in length) and fixed immediately in a solution containing 4% paraformaldehyde and 0.2% glutaraldehyde, diluted in Sorenson’s phosphate buffer 0.2M, pH 7.2, then placed in a graded sucrose series diluted in Sorenson’s phosphate buffer 0.1M as follows: 0.7M for 4h at 4°C, 1.5M for 4h at 4°C, and 2.3M sucrose overnight at 4°C. Then, samples were infiltrated with Jung Tissue Freezing Medium embedding matrix (Leica Instruments GmbH, Nussloch, Germany), overnight at 4°C. Transverse slices of 40 μm thickness were obtained with a cryostat at -20°C (Jung CM 1500, Leica instruments, GmbH, Nussloch, Germany). Sections were rinsed with PBS1x, and stained with aniline blue for 10 minutes, in darkness at room temperature. After incubation, the sections were rinsed with PBS and observed under a Zeiss Axio Observer Z1 microscope (Carl Zeiss GmbH, Munich, Germany) using an excitation filter at 405 nm and emission wavelength at 435-490 nm. To set the exposure time, infected samples from both lines were used to define the optimal condition to avoid overexposure. For both lines, at least four non-serial sections from three different biological replicates were analyzed. Unstained sections were observed with the same settings used for the stained ones, as controls. Image analysis was performed to evaluate the callose deposits. The number, intensity, and integrated density (the ratio between number and intensity) of callose deposits were quantified as previously reported (Zavaliev and Epel, 2015) using FIJI software (Schindelin et al., 2012) and the ReniyEntropy algorithm.

### Statistical analysis

Statistical analyses of all the results were performed using R with Rstudio software Version 1.1.456 (RStudio Team (2020). RStudio: Integrated Development for R. RStudio, PBC, Boston, MA). To identify significant differences between the proportions of damaged and undamaged SEs, the χ2 test was used with p<0.05. For the other results, a one-way ANOVA was performed, followed by a post-hoc test to identify significant differences between the plant lines. Conformity to the normal distribution and homogeneity of variances were checked with Shapiro-Wilk’s and Bartlett’s tests, respectively. Where necessary, data were normalized with a Box-Cox transformation. The non-parametric Wilcoxon test was used to determine significant differences in the average number of vesicles between the lines (*wild type*, *syta-1*) with p < 0.05. For the area of SEs, the diameter of the vesicles, perimeter, and the kurtosis, the Bonferroni test was used to determine significant differences among the lines (*wild type*, *syta-1*) with p < 0.05. As for the ER damage in the *syta-1* line, the Student’s t-test was used to determine significant differences between the lines (*wild type*, *syta-1*) with p < 0.05. For evaluation of the callose deposition, two-way ANOVA with a post-hoc t-test was used to determine significant differences between healthy and infected plants with p < 0.05.

## Results

### SYTA localizes to the sieve elements

Previous GUS analysis revealed that *SYTA* is highly expressed in vascular tissues (Schapire et al., 2008). Our observations confirmed that the *SYTA* is widely expressed throughout the plant, with a particularly strong expression in vasculature and roots (Fig. 2A). To test if SYTA localizes to SEs, we crossed a *syta-1*;gSYTA-RFP line, where the genomic *SYTA* sequence is fused to RFP and expressed under the *SYTA* promoter, with a SUC2::GFP line, which expresses GFP under the *SUC2* promoter and serves as an identifier of SEs (Doidy et al., 2012). The *syta-1*;gSYTA-RFP plants exhibited clearly fluorescent signals within vascular tissues (Supplementary Fig. S1B). In the crossed line (*syta-1*;gSYTA-RFP x SUC2::GFP), co-localization of GFP and RFP signals was detected mainly in SEs as well as in some adjacent phloem cells (Fig. 2B). A similar expression pattern was obtained using CALs7, another SE identifier (Supplementary Fig. S1C).

**Figure 2.**
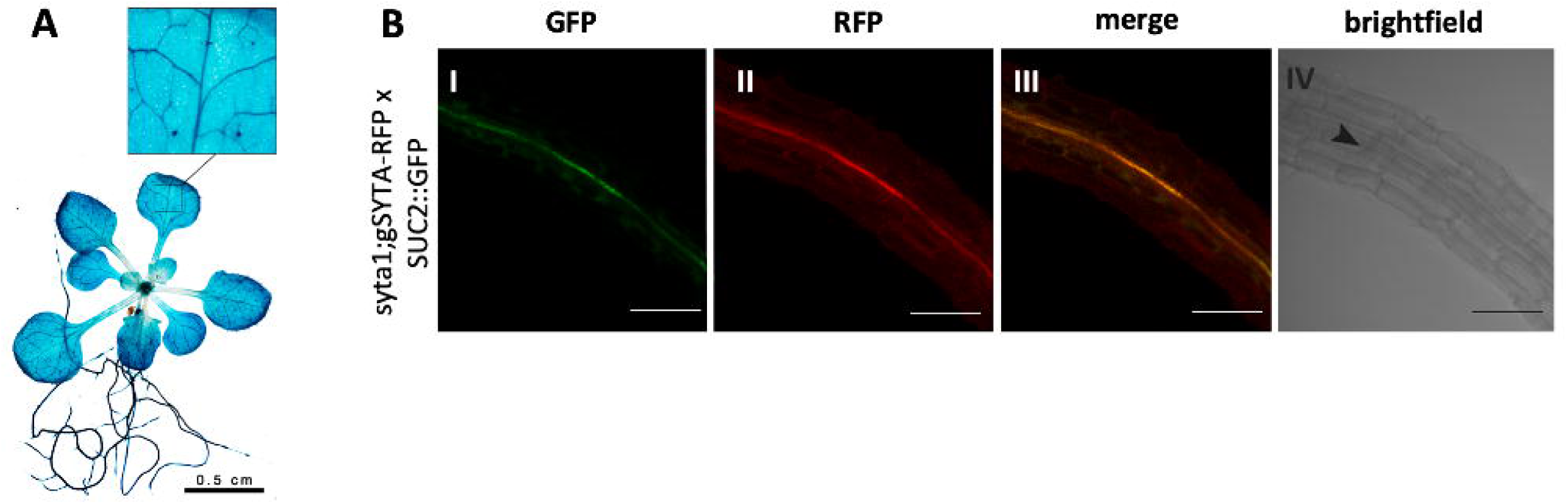
Localization of SYTA in Arabidopsis. (A) GUS assay in *Arabidopsis*, showing *SYTA* promoter expression in the leaf vasculature. Bar= 0.5 cm. (B) localization of *syta-1*;gSYTA-RFP x SUC2::GFP in root tips, showing SYTA localizes in the sieve elements. (B-I) Localization of the GFP signal; (B-II) Localization of the RFP signal; (B-III) merge of the two fluorescent signals; (B-IV) Brightfield micrograph of the sections observed. Arrows indicate the vasculature area. Bar = 100um.

### SE endomembrane system is damaged in *syta-1* mutants

To determine whether SYTA plays a role in the structure and the position of SE endomembrane system, we compared the morphology of SEs and enclosed SER/PM in *wild type* and *syta-1* mutant plants in TEM sections. In the *WT*, the SEs appeared intact, with the SER and plasma membrane appressed generally to the cell wall (Fig. 3A-E). The SER showed a highly regular architecture, with well-defined cisternae (Fig. 3A-E). In contrast, in *syta-1* plants, the endomembrane system did not appear in proximity with the SE wall (Fig. 3F-K). Although few SEs retained their normal morphology (Fig. 3F), the majority showed various degrees of inner membrane disruption (Fig. 3G-K). In some *syta-1* sieve elements, SER cisternae were aligned parallel to the SE wall (Fig. 3F i), while in others, the SER was oriented perpendicularly (rotated at 90°) (Fig. 3F ii). Quantitatively, membrane abnormalities (SER and PM not pushed against the SE wall) were observed in 19% of *wild type* SEs compared to 56% in *syta-1* mutants (Fig. 3J).

**Figure 3.**
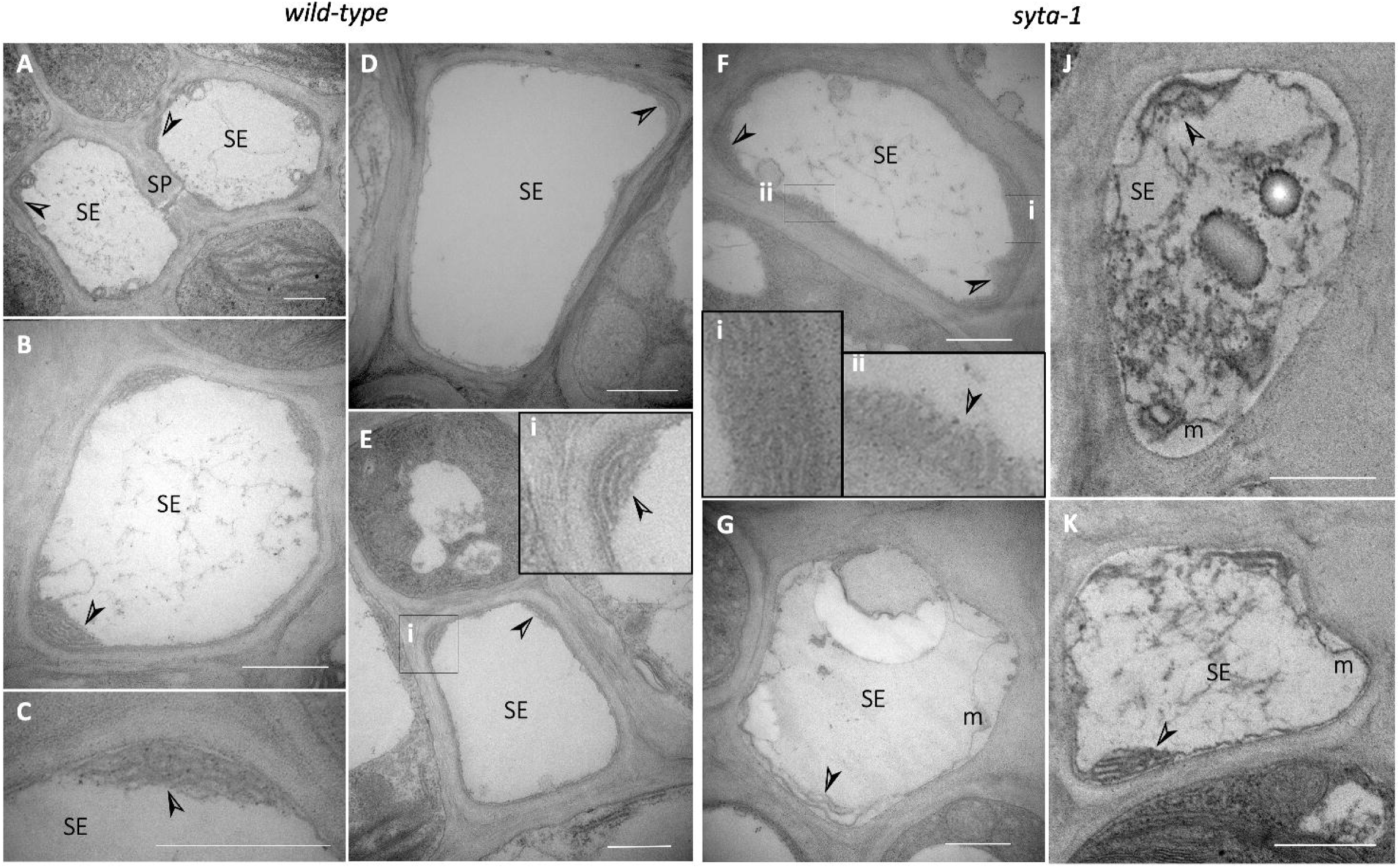
TEM micrographs of *wild type* and *syta-1* phloem from adult Arabidopsis leaves midveins. In *wild type* **(A-E)**, SER has a conventional cisternal structure. The SER is firmly appressed to the plasma membrane in all cells observed (A-E) and stacks and cisternae were well defined (B, C, inset E-i). The sieve plate appears as a defined channel, with a collar of callose lining the pore (A). In *syta-1* **(F-K)** SER is damaged (arrowhead in G, J, K) and distanced from the cell wall. The SER appears collapsed (Fi) and often display a 90-degree-rotation (Fii) in *syta-1* plants. The stacks are not well defined or organized in a compact shape (G-K). Damages to the inner membrane are frequent in *syta-1* plants: the whole membrane system appears distanced from the cell wall (G-K) and contains many vesicles without a regular shape (J). SE=sieve element, SP=sieve plate, CC= companion cell, arrowhead = endoplasmic reticulum, m=damaged membrane. Bar = 1 μm.

Because the lack of SYTA may interfere with SE development, we quantified and compared the areas of the SE lumina. On average, *syta-1* plants exhibited a significantly smaller SE area compared to *wild type* plants, with a value of 6.35±0.38 vs 8.25±0.39 (μm^2^) respectively (Fig. 4B). We then counted the number of membrane perturbations and vesicles per SE, as well as the average diameter of intact vesicles (Fig. 4C-D). The *syta-1* showed an increased number of vesicles and SER membrane and PM detachments relative to *wild type* (Fig. 4C). To test if the membrane integrity could have a role in the vesicle size, we measured the diameter of intact vesicles (Fig. 4D). The vesicle diameter in *syta-1* was significantly larger than in WT.

**Figure 4.**
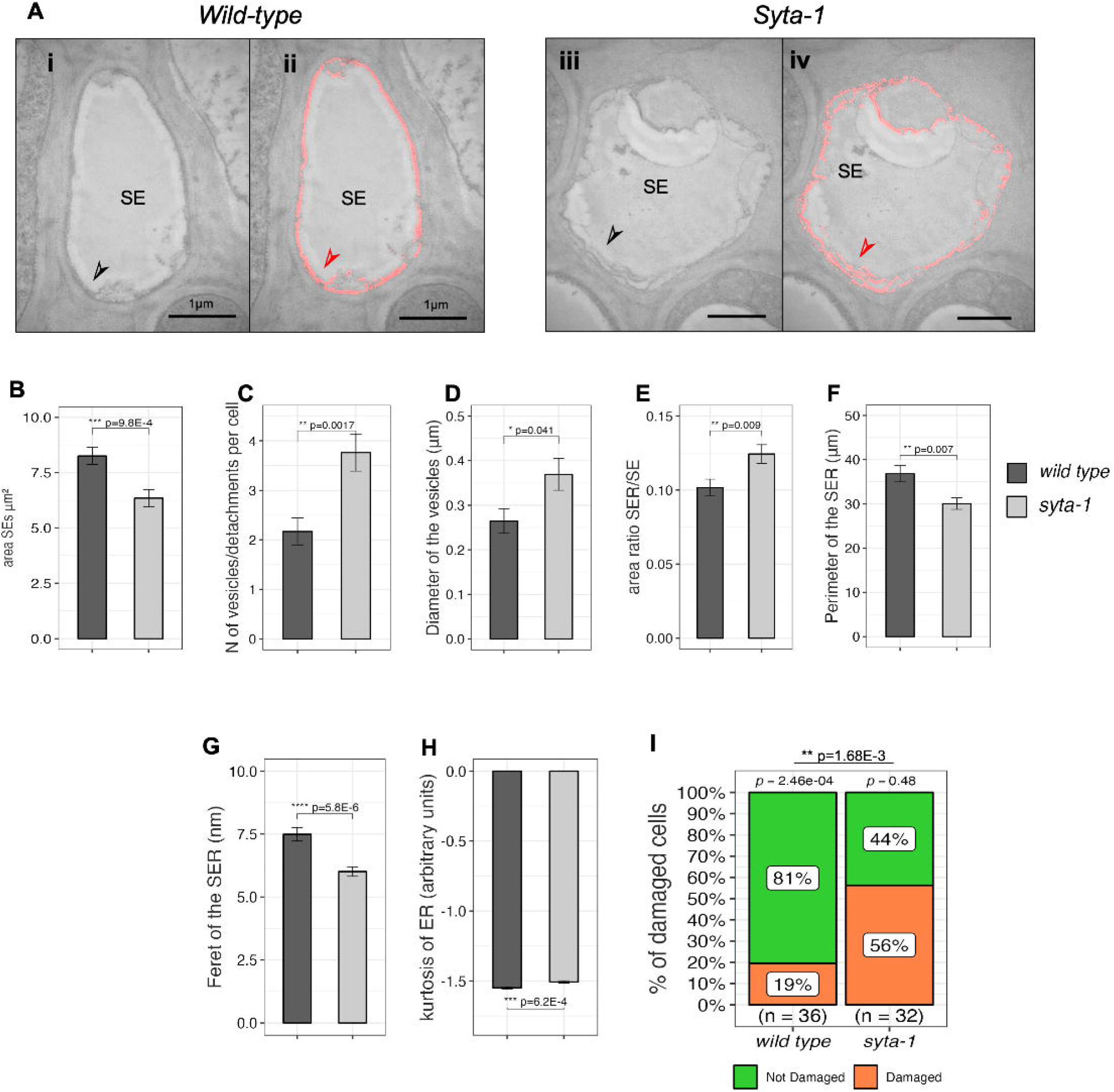
Quantification of SE membrane characteristics is *wild type* and *syta-1* sieve elements. (A) Representative micrograph of the SER/PM in *wild type* (i) and *syta-1* (iii) and overlay of the micrograph with the Ilastik mask (red pixels) (ii, iv). SE=sieve element, arrowhead = endoplasmic reticulum (SER/PM), orange portions = masked area. **(B-H) ER/PM segmentation in *wild type* and *syta-1***. (B) Mean area of the sieve-element lumina in *wild type* and *syta-1* lines. (C) Number of vesicles and/or detachments of the PM from the cell wall and (D) diameter of well-formed vesicles in *wild type* and *syta-1* plants. (E-H) Shape indicators of the SER/PM in *wild type* and *syta-1* plants: area of the membrane/sieve-element lumen ratio (E), perimeter of the SER (F), Feret diameter of the SER (G) and kurtosis of the SER shape (H). (I) χ-test for membrane (SER and PM) damage incidence in *wild type* and *syta-1* lines. In the bar plots, data are expressed as mean value ± standard error of the mean from 3 biological replicates for each line. Asterisks show significant differences among the means * for a p<0.05, ** for a p<0.01, *** for a p<0.001, **** for a p<0.0001 (Wilcoxon test in C and Bonferroni test in D, Student’s t-test for B, E to H.

For a quantitative analysis of SER morphology, we used supervised machine learning to generate segmentation of the SER and PM (Fig. 1) in both *wild type* and *syta-1* (Fig. 4A). This segmentation enabled automatic quantification of structural membrane parameters, such as the total area, perimeter, Feret diameter, skewness, and kurtosis (Fig. 4E-H, Supplementary Fig. S2A-C). The analysis revealed that the SER/PM-to-lumen area ratio was significantly higher in *syta-1* plants (Fig. 4E). In contrast, the total membrane perimeter was significantly higher in the *wild type* plants (Fig. 4F), and the Feret diameter of the single particles was also greater in *wild type* compared to *syta-1* plants (Fig. 4G). Additionally, kurtosis of the membrane was significantly different between *wild type* and *syta-1* (Fig. 4H). Together, these results indicate that the structure of the membrane and of the SER specifically, in *syta-1* mutants was deformed in many ways.

### 14C export is inhibited in syt-a mutant plants

To determine whether changes in sieve element morphology affect carbon export, we measured the fixed ^14^C over time, up to 2500 minutes following ^14^C labeling in *wild type* and *syta-1* plants (Fig. 5A). Relative ^14^C fixed standardized values (stVs) were consistently higher in *syta-1* plants than in wild type, with statistically significant differences observed at 500-, 1250-, 2000- and 2750-minutes post inoculation (Fig. 5A). We then analyzed the carbon loss from the leaves by using an exponential decay function to describe the 14C loss in both *wild type* and *syta-1* plants (Fig. 5B-C). In *wild type* plants, the curve fitted with an exponential decay phase started 500 minutes after the labeling, whereas in *syta-1*, this phase was delayed until 900 minutes (Fig. 5B-C). These findings suggest that ^14^C export is more efficient in *wild type* plants and is impaired in *syta-1* mutant.

**Figure 5.**
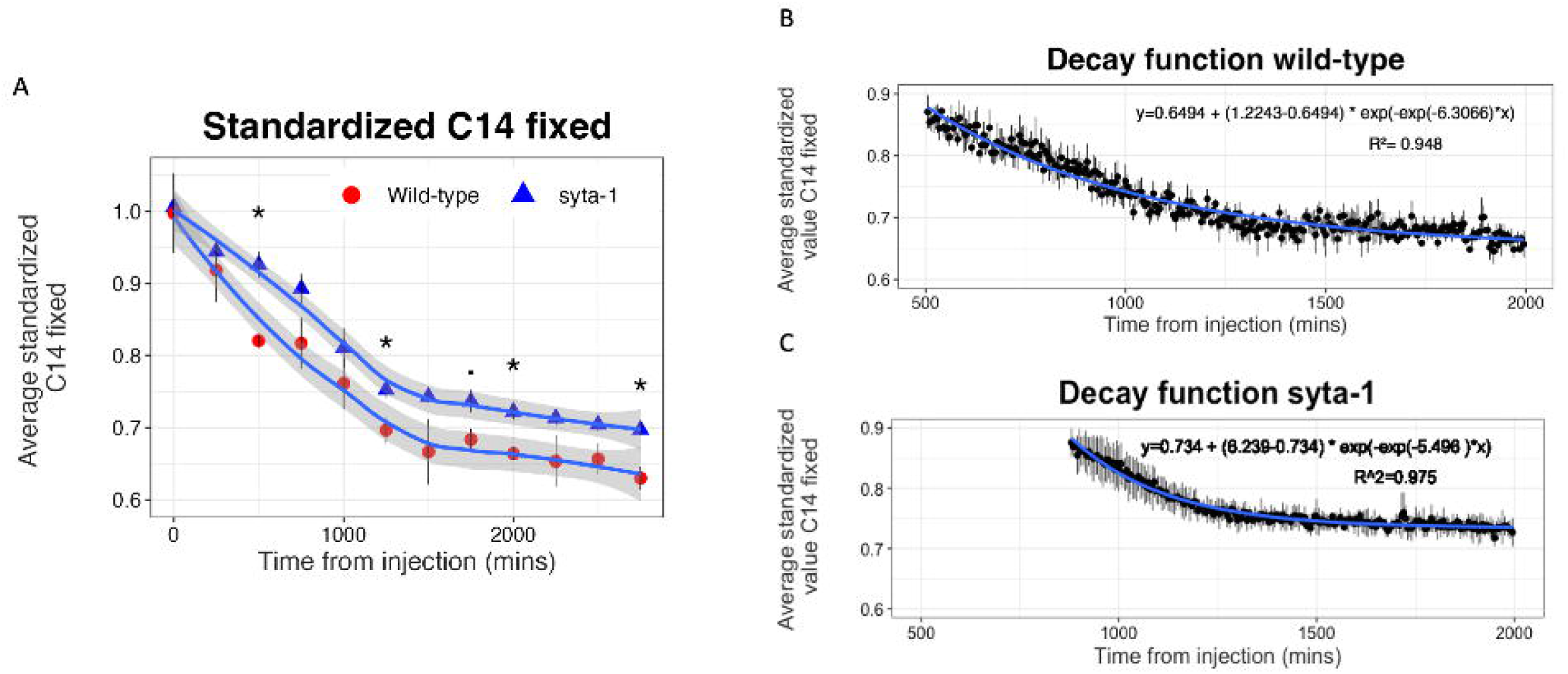
^14^C export it *wild type* and *syta-1 plants*. Standardized exportation value along the experiment (A) and decay functions (B-C). (A) Loss of ^14^C-labeled compounds from rosettes in *wild type* and *syta-1* plants during the export period. Single points express the average value ± SEM of three biological replicates. Significant differences among the means are reported as ° for a p<0.1, * for a p<0.05 according to the t-test. Decay functions of the isotope in the exponential phase in *wild type* (B) and *syta-1* (C) plants. Single points express the average value ± SEM of 3 biological replicates.

### *AtSYTA* downregulation reduce callose deposition in phytoplasma-infected plants

To investigate a role of SYTA in response to phloem-limited pathogens, we infected Arabidopsis *wild type* and *syta-1* knockdown plants with CY-phytoplasmas. 21 days after the end of the inoculation access period (IAP), both lines showed typical phytoplasma-associated symptoms, including yellowing, phyllody, and petiole elongation (Fig. 6A). Despite symptom development, fresh weight did not differ between healthy and infected plants of either line (Supplementary Fig. S3). However, the SPAD index, which reflects chlorophyll content, was lower in infected *syta-1* line compared to healthy *syta-1* and healthy *wild type* (Fig. 6B).

**Figure 6.**
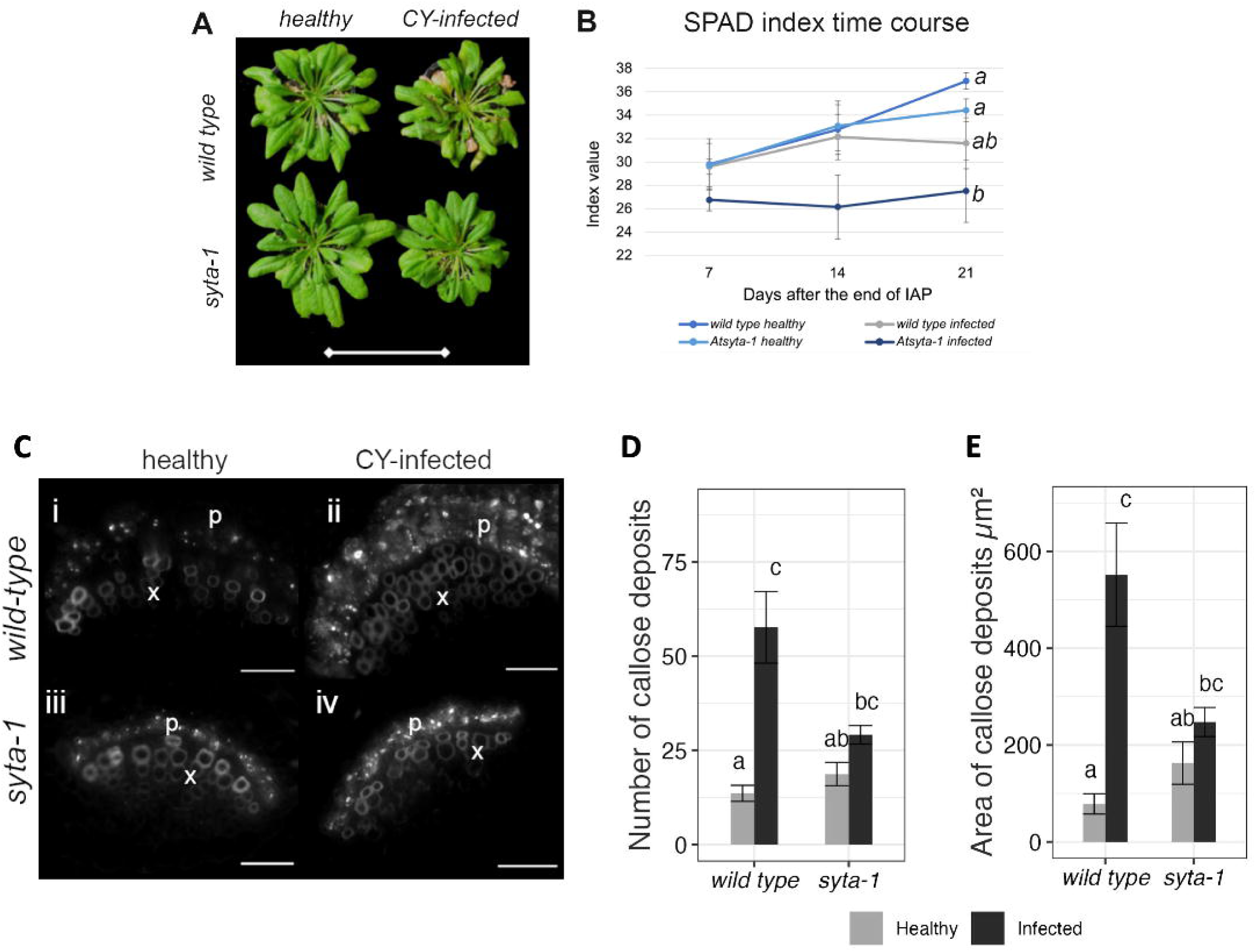
Biometric evaluations and callose imaging analysis on healthy and CY-infected *wild type* and *syta-1* plants. (A) The morphology of both lines appears similar among healthy samples belonging to the two lines. Infected plants seem to be slightly smaller than healthy ones. Bar is 10 cm. (B) SPAD (soil and plant analysis development) measurement for the chlorophyll content shows a lower index value for infected mutant line 21 after the end of IAP. The scatterplot expresses the average ±SE (7 biological replicates). Statistical analysis was carried out at the third measurement point. The barplot expresses mean±SE. Different letters express differences between the means according to two-way ANOVA followed by Tukey’s test as post hoc test with p<0.05. (C) Representative micrographs of healthy (i-iii) and infected (ii-iv) *wild type* (i-ii) and *syta-1* (iii-iv) stained with aniline blue. Fluorescent dots indicate callose deposits in the sieve plates. X = xylem, p=phloem, Bar = 200 um. Count (D) and area (E) of the callose deposits. Data are expressed as mean value ± standard error of 3 biological replicates. Different letters express significant differences between the means (Tukey’s test with p<0.05).

DAPI staining revealed no significant difference in phytoplasma titer between *wild type* and *syta-1* plants (Supplementary Fig. S4A-D). However, when we examined infected phloem using TEM, we observed some structural distinctions (Supplementary Fig. S5A-B). Interestingly, the typical funnel-like structures between SER and phytoplasmas were also present in both wild type and *syta-1*, which indicates that SYTA is not required to establish this mode of interaction (Supplementary Fig. S6).

To reach a better impression of callose accumulation, we stained healthy and infected *wild type* and *syta-1* samples with aniline blue (Fig. 6C). In *wild type* plants, infection triggered more callose accumulation compared to healthy controls (Fig. 6C i, ii, Fig. 6D). However, in *syta-1* plants, infection did not lead to callose accumulation, and callose levels stayed unchanged from uninfected samples (Fig. 6C iii, iv; Fig. 6D). This was corroborated by the quantitative data, which showed a 68% increase in callose deposit area in infected *wild type* plants, while no significant change was seen in *syta-1* line (Fig. 6E). Collectively, the results highlight the role of SYTA in regulating SER integrity and activating callose-based defenses during phytoplasma infection, possibly through modulation of ER–pathogen interactions at MCSs and physically retarding the movement of the pathogen.

## DISCUSSION

### Partial loss of membrane integrity in *syta-1* mutants

The SE endomembrane system was more damaged in *syta-1* mutants than in wild-type plants (Fig. 4I). One cause of SE membrane distortion could be a loss of membrane integrity (Schapire et al., 2008). Although not directly studied here, we observed an increased number and diameter of vesicles (Fig. 4C-D) and an increased number of undefined SER cisternae in the *syta-1* plants (Fig. 3 and 4A). Moreover, the higher number of damaged membranes in *syta-1* mutants (Fig. 4I) points to the same explanation. SYTA can restore and maintain the membrane integrity and support resealing (Schapire et al., 2009, 2008; Yamazaki et al., 2008). Hence, its absence could cause breakage of membranes due to the inability of reparations independently from the nature of the membrane (either ER or PM). Abiotic stresses underlined the role of SYTA in membrane maintenance of integrity and stability. Osmotic or mechanical stress causes membrane disintegration, and the damage is exacerbated in plants with downregulated SYTA (Balsam and Bush, 2022; García-Hernández et al., 2025; Krausko et al., 2022; Ruiz-Lopez et al., 2021; Schapire et al., 2008). In fact, among the pathways compromised by an abiotic stress in syta mutants are those directly involved in the maintenance of the PM stability (Kusá et al., 2025).

In this study, *syta-1* plant SE PM undergoes plasmolysis even without any other stress (Fig. 3 G-K), a phenomenon that we observed sometimes in parenchymatic cells as well (data not shown). This may result from a synergic action of the lack of anchoring points or membrane stability and the lack of adequate Ca^2+^ signaling related to SYTA reduction ((Juan Luis Benavente et al., 2021). It is well established that the Ca2+ can be involved in the plasmolysis and the adhesion of the PM to the CW (Hayashi et al., 2006). Additionally, the plasmolysis phenomenon may worsen the ER damage by acting indirectly on its motility: although the ER remains adhered to the PM, it appears less motile and with a different propensity for the partitioning (Cheng et al., 2017).

### In syta-1 mutants, SER cisternae are more fragmented and separated from cell wall than in *wild type* plants

Image segmentation revealed that the SE membranes appeared to be somehow damaged. Given that the ER constitutes the majority of the membranes inside the SE and that it covers the most aggregate surface in a 2d image, we infer that also the majority of the damage described hereafter occurs at the level of the ER. SE membranes showed a lower average aggregate perimeter in *syta-1* mutants, a lower Feret diameter and a higher value of kurtosis than in *wild type* plants (Fig. 4F-H). By contrast, the number of free vesicles (Fig. 4C), which are generally larger Fig. 4B), is higher in *syt1* mutants. Perimeter describes the organization of the membranes along the sieve tube edge: a higher value of the perimeter corresponds to a distribution of SER and PM around the edge of the sieve element, in contrast, lower values indicate that the membrane collapses and hypothetically accumulates on one side of the cell (Figure 4, model proposed in Figure 8). Larger aggregate SER perimeters coincide with a regular ER organization having an appressed position against the SE PM along the SE border (Fig. 4F). The smaller Feret diameter, which expresses the average radius of the particles (Fig. 4G) evidences larger ER fragmentation in *syta-1* plants (Fig. 4G). Kurtosis is a measure of the flatness of SER cisternae thus, the higher kurtosis values of the *syta-1* SER indicate a less reticulated shape as compared to the *wild type* (Fig. 4H). The results collectively demonstrate that, in *syta-1* mutants, the SER membranes separate from the PM, the SERs have a less discrete configuration, and SEs contain more intracellular and larger vesicles than in *wild type* plants. Altogether, the results infer that SYTA is responsible for cisternal SER anchoring and SER tethering to the PM, coherence and shape of the SER and maintenance of membrane integrity. Another reason for SER abnormalities might be a reduced ability to a proper development and organization of new membranes. Because the ER is a highly dynamic organelle, a continuous membrane degradation and reconstruction are constantly required (Li et al., 2025a). We did not observe any difference in the area of the SER when *SYTA* is downregulated, while the relative area (express as area of SER/area of SE) was lower in *syta-1* (Supplementary Fig. S2A; Fig. 4E). suggesting that SYTA does not play a role in the formation of new SER but rather its anchoring and the organization of its shape. A well-organized ER in stacked karmellae or lamellae (as is the case in the SER) is designated as an organized smooth endoplasmic reticulum (OSER) (Sandor et al., 2021). Since membrane-membrane connections are essential for OSER formation (Sandor et al., 2021), the lack of SYTA could be an obstacle for proper cisternal arrangement in *syta-1* mutants.

After gentle tissue fixation for EM, Ehlers and coauthors demonstrated the presence of ER-PM clamps SEs, apparently involved in maintaining the immobility of the SER (Ehlers et al., 2000). Given a lack of tethering ability in *syta-1* plants (Fig. 3F-K), we hypothesize that SYTA may reside in these clamps in order to contribute to the SER stability as inferred by previous studies. In cells that do not belong to the phloem, the lack of SYTA destabilizes ERs and the ER tubules appear more dynamic compared to *wild type* plants (Ishikawa et al., 2018; Levy et al., 2015; Siao et al., 2016). Interestingly, downregulation of *AtSYTA* can negatively impact the stability and abundance of VAMP-associated EPCSs (V-EPCSs), while the absence of VAP does not affect the formation of SYTA-associated EPCSs (S-EPCSs) (Siao et al., 2016).

### Carbon export is suppressed in *syta-1* mutants

Although the decline of a carbon tracer signal is often interpreted as leaf C export, respiration also plays a role in the loss of photosynthates from the leaf (Kölling et al, 2013). Thus, without measuring respiration, it is not possible to distinguish which portions of the decline of the 14C signal result from export or respiration. However, reduced export is likely to increase respiration, tending towards hiding, rather than accentuating differences in export. This is because respiration responds to carbon supply in the form of starch reserves to maximize starch utilization without completely depleting reserves (Suplice et al. 2014; Stitt and Zeeman, 2012). Thus, leaves with lesser capacity for export will metabolize a greater portion of their C within the leaf, which make differences in export more difficult to detect. Thus, the differences in actual C export from rosettes between syta and *wild type* are likely to be greater but not lesser than those measured by the decline of ^14^C signal.

Our study shows that ^14^C export in syta-1 plants is slower than that in *wild type* plants (Fig. 5). According to the current state of knowledge, mass flow through sieve tubes follows the fluid-dynamic principles (Thompson and Holbrook, 2003). Consequently, new formulae to describe phloem transport were developed (Minchin and Lacointe, 2017; Pickard, 2012), which include any hindrance to proper mass flow. According to Poiseuille’s equation, any modification in the morphology of the sieve tube can affect its conductivity, assuming a low Reyonolds number (Jensen et al., 2016). For instance, sieve-tube diameter and the diameter and density of sieve pores were proposed to be the most critical parameters in determining the mass flow rates (Mullendore et al., 2010).

In *syt1-a* mutants, several determinants of mass flow rate are affected. First of all, the cross-sectional area of SEs in *syta-1* plants was lower than in *wild type* plants (Fig. 4B), leading to a decreased mass flow rate (Thompson and Holbrook, 2003). In addition, the irregular arrangement of SER cisternae in mutants could be a source of friction that retards mass flow. Furthermore, sieve-pore dimensions controlling mass flow rates are affected. The *syta-1* mutants showed an increased callose deposition around the sieve pores when compared to the *wild type* (Fig. 6 C-E). As demonstrated previously, callose deposition at the sieve pores can slow down the phloem transport speed (Bernardini et al., 2022a). And as a last note in this frame, loose ER vesicles in mutant plants could amass against the sieve plates which could be another source of mass flow resistance. Lastly, it is also possible that the function of SYTA in other parts of the plants can indeirectly contribute to this difference.

### Failing membrane tethering implicates phloem infections

Because SYTA was shown to decrease the infection rate of different plant viruses, among them are also phloem-limited viruses (Uchiyama et al., 2014), we studied the interaction between Arabidopsis and phloem limited CY-phytoplasma. While a difference in phytoplasma titer between *wild type* and *syta-1* plants was not observed (Supplementary Fig. S4), the downregulation of *SYTA* resulted in reduced disease symptoms (Figure 6). It is possible that SER is not required for phytoplasma replication and/or movement, or the pathogen successfully reproduces in those SEs having an intact SER, which yet occurs in 44% of the SEs in *syta-1* mutants (Fig. 4I). However we strongly believe that the damage of the inner membrane (either PM or ER) inside the sieve element could somehow influence the movement of the phytoplasmas: phytoplasmas were suggested to exploit the actin for their movement (Boonrod et al., 2012; Buxa et al., 2015; Galetto et al., 2011), and it is possible that the reduced amount of SYTA could impact the actin binding as well. The observed colocalization of actin with SYTA could enforce this hypothesis (Levy et al., 2015). On the other hand, the presence of the cytoskeleton in the SEs is still controversial and a matter of debate (Van Bel and Musetti, 2019). Over the years, some authors have strongly asserted the presence of the cytoskeleton (Hafke et al., 2013), while others have rejected this hypothesis, demonstrating that cytoskeleton components are remnants from sieve element maturation (Knoblauch et al., 2018). Thus, the relation between phytoplasma and actin remains elusive.

Another reason for the reduced symptoms in *syta-1* plants may be the impairment of the callose deposition in infected sieve elements. Interestingly, in *wild type* plants, the downregulation of *SYTA* resulted in slightly higher callose levels in the phloem; however, after infection, we could see a strong increase in callose levels in the *wild type* plants, while in *syta-1* the basal callose levels remained stable and did not increase. It was shown recently that upon perception of microbe-associated molecular patterns (MAMPs), SYTA rapidly accumulates to PDs and recruits a putative calcium-permeable transporter, ANN4, to promote a localized, PD-associated Ca^2+^ elevation, leading to callose-dependent PD closure (Li et al., 2025). Callose constriction of sieve pores to obstruct pathogen spread is an important mechanical defense response to phloem pathogens (Bernardini et al., 2024, 2022a). However, callose accumulation in the phloem can also restrict sugar transport and lead to increased disease symptoms. Interestingly, and Calcium is primarily stored inside the ER (Pirayesh et al., 2021), and previous studies have speculated about its role in callose deposition (Furch et al., 2007; Hafke et al., 2009; van Bel et al., 2014). The question arises if a similar mechanism holds for SER, which is the only storage organelle in SEs (Oparka and Turgeon, 1999). Reduced callose deposition in *syta-1* might reflect a reduced release of Ca^2+^ from the storage sites so that the threshold for callose synthesis is not attained. Therefore, the callose pattern we observed in *syta*-1, with higher basal levels that do not increase after infection, could explain both the partial limitation of the pathogen leading to its unequal distribution, and the reduced symptoms, similarly to what was also found with the phloem-limited bacteria *Candidatus* Liberibacter asiaticus in citrus (Sarkar et al., 2024) Phloem-restricted pathogens may have adopted evolutionary solutions to establish trophic relations with the SER. Phytoplasmas have shown to possess attachments to the SER and the PM (Buxa et al., 2015; Park et al., 2021) and to disrupt SER homeostasis (Inaba et al., 2023; Musetti et al., 2023). Some evidence exists that CY-phytoplasmas are attached to the host’s SER through funnel-like structures. Such attachment attributes (Supplementary Fig. S5) might be part of a pathogen’s “feeding mechanism” to acquire substances from the ER (Pagliari et al., 2017, 2016; Rossi et al., 2018). In this scenario, the importance of SER tethering might be pivotal in forming this connection between the pathogen and ER also as a tool for phytoplasma replication.

### Conclusions

The phloem is the vascular tissue responsible for the transport of photosynthates and other organic molecules such as amino acids, phytohormones, and RNA, from source photosynthesizing leaves into sink tissues such as young leaves, flowers, fruits, and roots. Phloem endomembrane system is unique and still poorly understood while it plays a pivotal role in phloem biology, where the system is extremely simplified (Van Bel, 2003). Here, we describe a mutation of the gene encoding the *SYNAPTOTAGMIN A* in MCSs that disrupts inner sieve element membrane morphology by acting on the SER shape and the stability of the PM. SYTA also impacts physical aspects of phloem biology, such as the plugging of the phloem, the translocation from source to sink, and the establishment of the disease or the spread of diverse plant pathogens. Overall, our results show that the membrane contact sites play a key role in phloem functions. Closer examination of the SER tethering and pathogen interactions will provide new insights into the functions of the phloem and in understanding the way energy is distributed throughout the plants.

## Acknowledgment

We thank Donielle Turner, Chunxia Wang and Myrtho O’Pierre for technical support with the experiments. Seeds of the SUC2::GFP line were kindly provided by Professor Idan Efroni (Hebrew University of Jerusalem). The seeds of the CALS7::GFP line were provided by Dr Lothar Kalmbach (Sainsbury Laboratory, University of Cambridge). plasmid pCambia1301 +-1983, used for the GUS assay, was kindly provided by Prof. Sondra Lazarowitz (Cornell University). Support was provided by the National Institute of Food and Agriculture (grant 2020-70029-33197).

## Author contributions

**CB, RM, CV and AL** designed the research; **CB**, **SW** and **MK** performed of the research; **CB**, **CW**, **CV**, **MK**, **AVB** and **AL** analized the data; **CB** and **AL** wrote the manuscript

